# Honey bee egg composition changes seasonally and after acute maternal virus infection

**DOI:** 10.1101/2024.09.05.611496

**Authors:** Abigail Chapman, Alison McAfee, L. C. C Kenzie, Armando Alcazar Magaña, David R. Tarpy, Julia D. Fine, Zoe Rempel, Kira Peters, Rob W. Currie, Shelley E. R. Hoover, Leonard J. Foster

## Abstract

Honey bee (*Apis mellifera*) colonies depend on the reproductive output of their queens, which in turn is contingent on the care provided by worker bees. Viral infections in queens can compromise reproductive output, while worker infections can inhibit the successful functioning of a colony and its ability to care for the queen. Transgenerational immune priming (TGIP) occurs when queens transfer immune-related compounds or immune elicitors to their offspring, enhancing the ability of subsequent generations to resist infections. These maternal effects on offspring could positively impact colony health and resilience to viral infections, but little is currently known about TGIP for viruses. In this study, we investigate how viral infections affect the proteomic composition of eggs laid by virus-challenged queens (injected with a mixture of black queen cell virus and deformed wing virus B), both in controlled experimental settings and natural field conditions. Our results showed that virus-challenged queens upregulated immune effectors in their eggs and ovaries. In contrast, naturally infected queens from field surveys did not; there were no significant differences in egg protein, lipid, or metabolite composition related to maternal viral load or ovary size. However, egg collection date strongly influenced egg composition, likely reflecting seasonal variations in pollen resources. These findings highlight that while viral infections can induce transgenerational effects on egg proteomes under short-term experimental conditions, such effects are less apparent in natural settings and can be overshadowed by seasonal and other ecological factors.

## Introduction

Honey bee (*Apis mellifera*) colonies rely solely on the reproductive output of their singular queen to sustain their population ^1^. The queen, in turn, depends on the worker bees in the colony to care for and feed her, as she is primarily fed glandular secretions from the hypopharyngeal glands (HPGs) of nurse bees (young female worker bees whose main task in the hive is to feed the queen and larvae) ^2^. The quality and content of the diet of nurse bees impact the development of their HPGs ^3^, which in turn directly affects the reproductive output (egg laying rate) of a queen ^4^. The nutrition available to a colony depends on the resources available in the environment, which vary seasonally as different flowering plants are in bloom in the surrounding ecology. Acquisition of those resources depends upon the colony possessing a population of workers strong enough to retrieve those resources, which reciprocally depends on the queen’s ability to produce that workforce and workers’ ability to rear larvae to adulthood. Colony dynamics, therefore, depend on both resource availability and the colony’s ability to capitalize on those resources.

The presence of various parasites and pathogens, particularly viruses, in the colony can interrupt this system. For example, viral infections in workers can significantly affect the health and shorten the average lifespan of the workforce ^5^. The queen is likely protected, to some extent, from exposure to infectious agents through some aspects of social immunity: infected workers are less likely to feed the queen ^6,7^ and given the age-based division of labour in a colony, the queen is mainly cared for by young workers who have not yet left the hive and interacted with the environment (where they may acquire new infections)^1^. But, despite this layer of protection, honey bee queens are still regularly infected by most or all the pathogens which have been found in honey bees, including the many viruses that are nearly ubiquitous through most honey bee populations ^8^. Exactly how and when queens obtain these infections is yet unclear, though, as Kevill *et al*. ^9^ found that the pathogen profile of a colony had not influenced the pathogen profile of their queen 24 days after introduction. When a queen does become infected, though, those viral infections can then be vertically transmitted to a queen’s offspring ^10–12^. The exact mechanisms of vertical transmission are unknown for most viruses, but it has been shown that deformed wing virus (DWV) adheres to the chorion (eggshell) and probably infects the larvae upon hatching ^13^, negatively impacting the health and potential viability of that offspring.

Viral infections can also directly impede the ability of a queen to sustain the population needed for a healthy colony. We have previously established that increased levels of viruses are associated with smaller ovaries and beekeeper-identified poor-quality honey bee queens in samples collected from the field ^14^ and that viral infections reduce the queen’s ability to lay eggs in the laboratory ^15^ ― effects that are likely exacerbated when nutrition is limited ^16^. How the molecular composition of eggs changes in response to maternal virus infection is still poorly understood.

The composition and size of honey bee eggs is plastic ― several recent studies have shown that honey bee queens predictably alter the size of their eggs in response to the size of the colony in which they reside ^17,18^, and that this change in size is accompanied by molecular changes ^19^. These changes are examples of maternal effects, where the phenotype of the offspring is influenced by the mother through altered deposition of maternal proteins or mRNA ^20^. Changes to eggs associated with maternal infection could conceivably be maladaptive, as in the case of a trade-off, whereby one might expect the resources intended for developing oocytes in the ovaries to be diverted towards an immune response. This has been demonstrated in several *Dipter*a species, where the eggs laid by immune-challenged individuals have reduced protein content (reviewed in Schwenke et al. ^21^). In honey bees, a trans-generational cost of maternal viral immune activation was observed by Leponiemi et al. ^22^, wherequeens were orally inoculated with either inactivated DWV or phosphate-buffered saline (PBS), and their pupae were then injected with either DWV or PBS to see if the offspring benefited from the priming of the queens. The authors found that, paradoxically, the control (PBS-injected) offspring of queens that had been primed had higher levels of deformed wings compared to the offspring of unprimed (PBS-fed) queens. This suggests that maternal viral immune activation has the potential to actually harm the ability of their progeny to withstand the same infection.

Conversely, there could also be adaptive changes after maternal immune challenge, that benefit the embryo (that, in honey bees, develops rapidly ∼72 hours after egg-laying). This is true in cases of transgenerational immune priming (TGIP), whereby parents transfer either immune effectors or immunological triggers to their offspring, helping them prepare for a potential pathological encounter. This form of adaptive immunity transfer has been observed in many invertebrates (reviewed by Tetreau et al. ^23^). The mechanism for the transfer of bacterial immunological triggers has been elucidated in honey bees, but it is not clear if this occurs similarly for viral triggers. For bacteria, the yolk protein vitellogenin binds and transports pathogen fragments containing pathogen-associated molecular patterns from queen to offspring (as well as horizontally from workers to the queen in glandular secretions), ^24,25^ conferring resistance to bacterial infection in the larvae ^26^. In contrast, the evidence for similar immune priming of offspring after maternal virus exposure in honey bees is less clear, appearing to be idiosyncratic and dependent on context, including the route of maternal exposure and genetics ^8,22,27^. Recent work by Amiri et al. ^28^ found that the transcriptome of eggs containing DWV or sacbrood virus (SBV), likely obtained from existing maternal infections, functionally differed (determined by GO enrichments) from eggs with no virus detected, although very few genes were differentially expressed. While viral TGIP in honey bees may not involve the transfer of immunological triggers using the same mechanism as for bacterial immune priming, if the queen can prime her offspring to resist viral infections in some contexts, we hypothesize that this may involve the transfer of an increased abundance of immune effectors or other alteration of the composition of eggs she lays, influencing embryonic or larval virus resistance downstream.

We previously found that when honey bee queens were injected with either infectious virus, UV-inactivated inoculum, or saline controls, the queens injected with infectious virus had smaller ovaries and were less likely to resume laying eggs than queens injected with saline ^15^. Meanwhile, the queens injected with inactivated virus had an intermediate ovary phenotype. The results of these experimental manipulations support our previous findings that queens collected from the field with smaller ovaries tend to be identified by beekeepers as “failing” and are associated with higher levels of virus ^14,29^. Given this connection between virus infection and reduced ovary size, we hypothesize that the proteomic composition of eggs laid by honey bee queens would reflect the infection status of their mothers.

Here we first investigated if the anatomical differences in honey bee queen ovary mass resulting from a viral immune challenge translated to differences in egg protein composition. To further investigate if these findings are replicated in a field setting and expand on the transcriptional work done by Amiri et al. ^28^, we then examined eggs sampled from a field survey to see if the protein, lipid, and metabolite composition differs as a result of naturally-occurring queen virus infections. Given that both reproduction and immune activation are dependent on adequate nutrition ^4,16,30,31^ and that nutritional availability varies greatly across the season, we collected these samples at two different timepoints.

## Results and Discussion

### Immune effectors are upregulated in eggs after experimental immune activation of queens

In a previously published experiment ^15^, we collected eggs and the ovaries from queens seven days after they had been immune-challenged with an injection of either 1) a combination of infectious black queen cell virus (BQCV) and DWV-B (live group), 2) a UV-inactivated version of the same inoculum (inactivated group) or, 3) saline (control group). To see if the anatomical differences in the ovaries we previously observed translate to differences in ovary and egg protein composition, here we report the results of proteomic analyses of the ovaries of all the queens after sacrifice (n=9 in each group) and of their eggs. For egg samples, we were only able to perform proteomics on samples collected from n=7 control queens and n=6 inactivated virus-injected queens, since not every queen began laying eggs again after injection. The live virus group was excluded, as only n=2 queens resumed laying eggs after injection.

Of the 5,296 protein groups we quantified (1% FDR, based on reverse sequence hits) in the ovaries, 16 protein groups were differentially expressed in live virus-injected queens compared to the control queens (5% FDR, Benjamini-Hochberg correction, Figure 1a). Five of these were immune-related proteins (IRP30, transferrin, hymenoptaecin, beta-1,3-glucan binding protein, and carboxyesterase) that we also previously reported to be upregulated in the hemolymph of these queens ^15^. For the queens injected with inactivated virus, only one protein was differentially expressed (5% FDR) in the ovaries: the immune effector IRP30 (Figure 1b). In the eggs of the inactivated group queens, only two proteins were differentially expressed relative to the control group: IRP30 and another immune effector, transferrin (Figure 1c). These two proteins were also among the top proteins we previously reported as being upregulated in the hemolymph of these queens, among various other immune effectors ^15^. IRP30 is a uniquely hymenopteran, multi-functional protein that is upregulated in response to viral, bacterial, and fungal immune challenges ^32–36^ while also being expressed more highly in reproductively active, egg-laying individuals in both honey bees and bumble bees^33^. Transferrin is a protein associated with innate immunity across many organisms, including both insects and mammals and functions to sequester iron away from invading pathogens ^37^.

**Figure 1.**
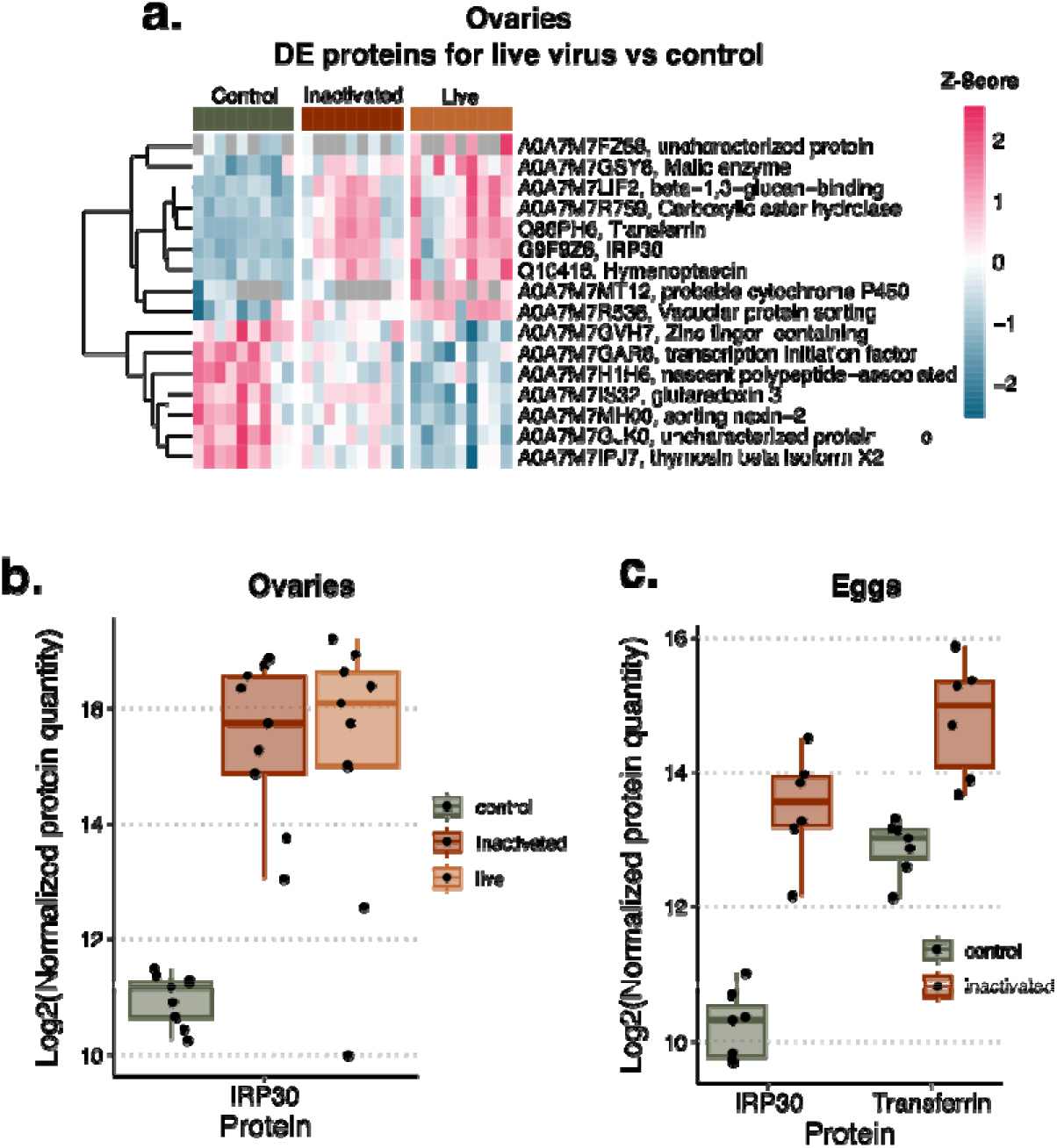
Immune proteins are upregulated in both the ovaries and eggs of queens injected with inactivated and infectious virus. For all differential expression analyses, the false-discovery rate was controlled to 5% using the Benjamini-Hochberg method. a) Sixteen proteins were differentially expressed in the ovaries of queens injected with infectious virus, compared to control queens (*n* = 9 for each group). Queens injected with inactivated virus are also shown (*n* = 9), but only IRP30 (bolded) was significantly differentially expressed in this group. Each row represents a protein showing the Uniprot accession number, and each column represents a sample. Proteins are clustered via Euclidean distance. b) IRP30 was the only protein significantly upregulated in the ovaries of queens injected with inactivated virus compared to the control queens (*n* = 9 for each group). c) IRP30 and transferrin were the only two proteins upregulated in the eggs of queens injected with inactivated virus (*n* = 6) compared to the control queens (*n* = 7).

Given that the proteomic changes between the ovaries and eggs of the queens injected with inactivated virus were nearly identical, and that much of the mass of the ovaries is made up of developing oocytes, one might infer that the differentially expressed proteins we observed in the ovaries of the live virus injected queens reasonably mirror changes to the proteins expressed in the eggs they laid. The increased differentially expressed proteins in the ovaries of the live virus-injected queens relative to the inactivated virus-injected queens support a previous suggestion by McMenamin et al. ^38^ that an interaction between the virus and host is likely responsible for eliciting a complete immune response. These authors observed that injection with dsRNA elicited only part of the immune response compared to injection with active virus. This effect also mirrors the intermediate phenotype we previously observed, where queens injected with inactivated virus had marginally smaller ovaries relative to the control queens ^15^.

### Egg composition is unchanged among queens with differing levels of natural virus infections

Our cage-based experiment demonstrated the maternal effects of an acute immune challenge with a combination of two common virus infections. To investigate how the effects of natural maternal virus infections compare, we collected queens and eggs from colonies managed by three beekeepers in BC and Alberta. The ovary mass and total virus copies (the logarithm-transformed sum of the RNA copy numbers of DWV-A, DWV-B, SBV, and BQCV) of these queens varied both within and among beekeepers ― across 83 queens collected, both ovary mass and total virus copies varied three-fold (Figure 2a). Surprisingly, we found no correlation between these two metrics (simple linear model including beekeeper and total virus copies as fixed effects with ovary mass as the response variable; F = 0.4855, p = 0.488, df_beekeeper_ = 2, df_total virus_ = 1), despite previously finding that ovary mass is negatively correlated with total virus copies in queens collected from the field ^14^. Given that potential maternal effects in the form of altered egg composition could extend beyond changes to the proteome, we performed lipidomics and metabolomics in addition to proteomics on the eggs from these queens. However, we found no differences (5% FDR, Benjamini-Hochberg correction) in egg proteins (5,574 detected), lipids (367 detected), or metabolites (997 detected) associated with maternal ovary mass or total virus copies. This lack of relationship is further reflected by no discernable clustering of samples by principal component analysis (Figures 2b, 2c, & 2d).

**Figure 2.**
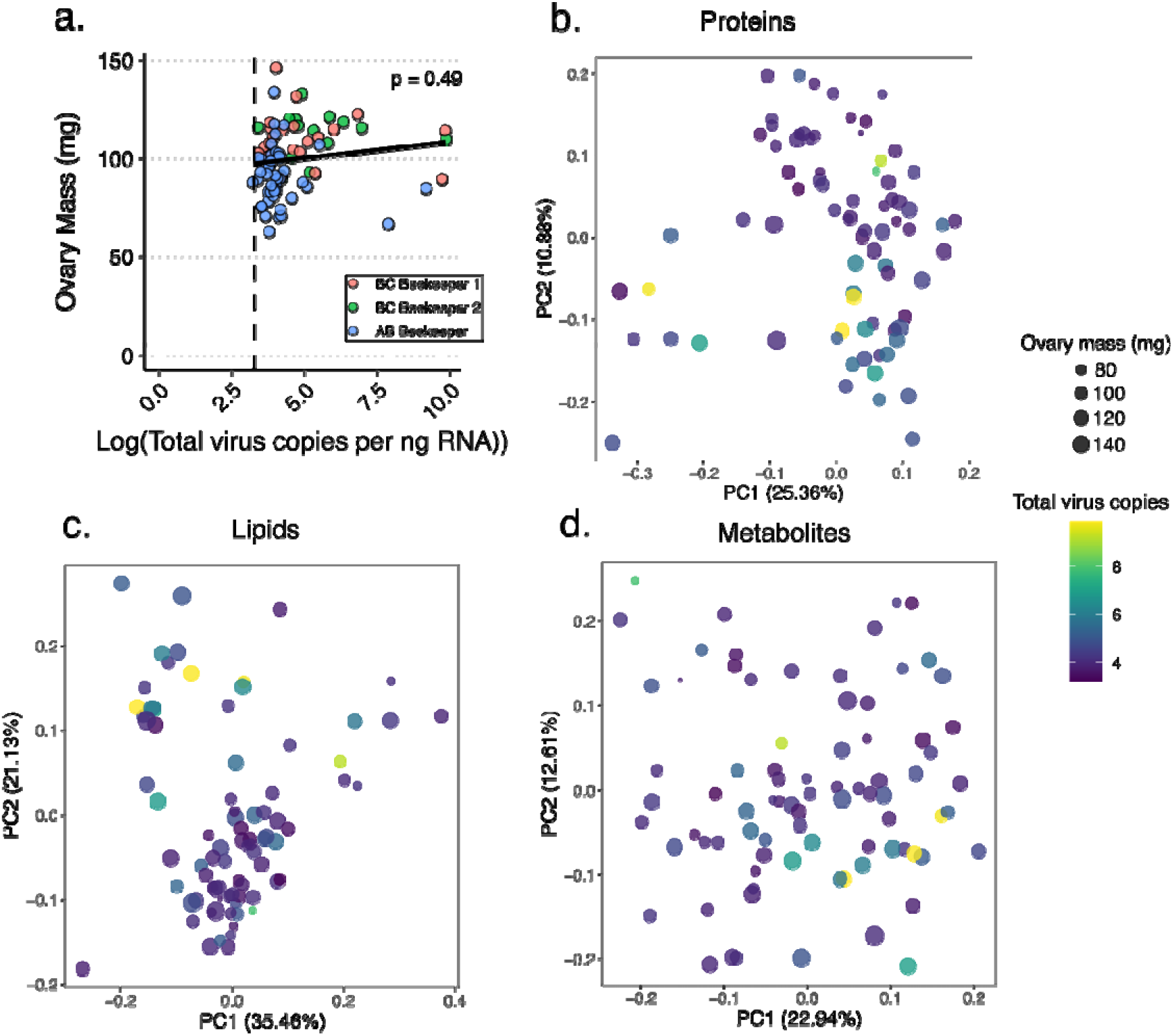
Variation among queens collected from the field. a) There was a large variation but no significant relationship between ovary mass and the total virus RNA copies in the ovaries of queens (F = 0.4855, p = 0.488, *n* = 87) collected from three beekeepers in British Columbia and Alberta. b) Proteomics analysis quantified 5,574 unique protein groups, but none were differentially expressed with respect to ovary mass or virus abundance (5% FDR, Benjamini-Hochberg correction). c) Similarly, lipidomic analysis identified 367 unique compounds in positive mode, but none were differentially expressed with respect to the same variables. d) The same lack of relationship with ovary mass and virus abundance was observed for metabolomics data, for which 977 unique compounds were identified across both positive and negative ion modes.

While we only collected eggs standing upright in each cell, which is indicative of newly laid eggs, the age of the eggs collected likely still varied considerably. Over the 72 hours of embryonic development between when an egg is laid and hatches into a larva, significant proteome changes occur ^39,40^ which may have partially obscured our ability to detect maternal effects. We also cannot rule out the possibility that some eggs laid by queens in these colonies differ in their protein, lipid, or metabolite profile but were removed by workers by some external quality control mechanism. Honey bees demonstrate an ability to detect infected and immune-stimulated nestmates across all life stages from larvae to adults, and remove the individual from the hive in a form of social immunity ^41,42^. Honey bees can also detect and remove diploid drone larvae^43^. Similar behaviour could have been at play here, where eggs exhibiting some altered signal were removed. The consistency in egg composition observed here could instead indicate the presence of some protective “quality control” mechanisms preserving the composition of eggs that queens lay despite changes to ovary activation and immune status due to that fact the transcriptome of early embryonic development is stabilized by homeostatic feedback loops ^28,44^. But, the lack of relationship between maternal virus level and ovary mass in this dataset indicates that perhaps the potentially chronic viral infections in this collection of queens are mild enough (the viral copies present in the field-collected queens were generally lower than the viral copies in all groups of the cage experiment) that they are not impacting the queen’s reproductive output or stimulating a transgenerational response. Given that we did see an effect of immune activation on the protein composition of ovaries and eggs in the laboratory cage experiment, this suggests that either 1) the effects of immune activation are magnified in a laboratory setting, especially given that the nutrition provided to a queen in a cage is likely reduced relative to a queen in a full colony environment, or 2) maternal effects are present in eggs only in the early phase of immune activation, whereas queens with infections in the field have reached some internal equilibrium where the activity of their immune system is no longer elevated.

### The lipidome and proteome of eggs collected from the field are strongly influenced by season

While we detected no changes in the eggs related to queen health metrics, there was a strong effect of the date of collection on the protein, lipid, and metabolite composition of the eggs collected from the two BC beekeepers on two different dates in mid-May and mid-June (n = 37 for each date). We found 691 protein groups (Figure 3a), 220 lipids (of which 34 are annotated, Figure 3b), and 126 metabolites (of which 18 are annotated, Figure 3c), to be differentially expressed regarding the date of collection.

**Figure 3.**
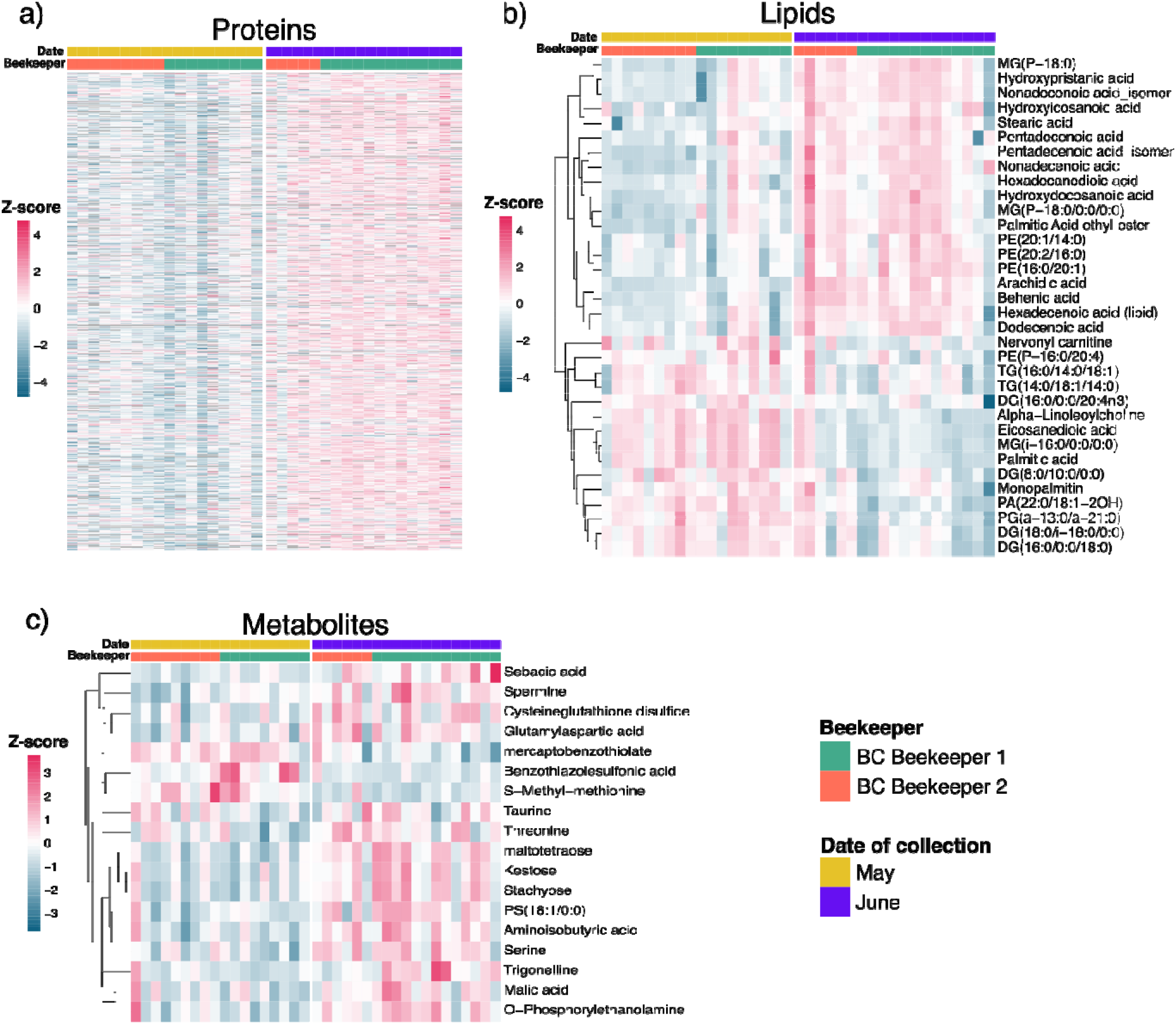
The composition of eggs changes with the season. The results of differential expression analysis were controlled to 5% FDR using the Benjamini-Hochberg method. Each row represents a protein or compound with rows clustered via Euclidean distance, and each column represents a sample. a) The 691 protein groups that were differentially expressed between May and June out of 5,574 identified protein groups. b) The 34 annotated lipids (from a total of 220 differentially expressed compounds) out of 367 total identified compounds, with differential abundance between May and June. c) The 18 annotated metabolites (from a total of 126 differentially expressed compounds) out of 997 compounds identified, with differential abundance between May and June.

Honey bees are known to exhibit seasonal variations in protein and gene expression, although most studies documenting this phenomenon compare extreme phenotypes (e.g., summer bees to winter bees), which experience very different life histories ^45,46^. Even though the samples taken in this study are temporally more proximate, one might still reasonably expect the changing environment over a single season to result in proteomic changes. It was surprising, though, that almost all these differentially expressed proteins were upregulated. We found that the gene ontology terms “large ribosomal subunit,” “unfolded protein binding,” “ribosomal subunit,” and “calcium ion binding” were significantly enriched (adjusted p < 0.05) in the proteins that were upregulated later in the season. These GO terms are related to some of the functional groups Han et al. found to be enriched in large eggs compared to smaller eggs including “Structural constituent of ribosome” and “Protein binding” ^18^. There were no GO terms significantly enriched in the proteins which were downregulated later in the season.

The 18 annotated metabolites that were significantly different were all upregulated at the second time point (Figure 3c). While the roles that most of these compounds play in honey bee eggs have not been previously described, some are involved in the same pathways that Han et al. found to be enriched in larger eggs ^18^, similar to the protein changes we observed. These authors also noted distinct changes in egg metabolites and proteins associated with the sampling date, but the precise temporal effect is unclear. There is evidence for many taxa, including mammals, amphibians, reptiles, and insects, that females shift to laying fewer, larger eggs later in the reproductive season ^47^. This contextualizes our finding that the proteins and metabolites we quantified as upregulated later in the season corresponded with similar markers in larger eggs.

Many of the annotated lipids that changed significantly over time in the eggs (Figure 3) are also found in pollen collected by honey bees ^48–50^. It is well-established that the protein and lipid content of collected pollen varies over the season ^49,51^ and that young worker bees convert the pollen they consume into nutrient-rich jelly to feed the queen and developing larvae ^52^. It is unsurprising, then, that the lipid composition of eggs varies seasonally as well, given the available resources to a colony.

## Conclusion

Following an experimental immune challenge with a combination of two common viruses (BQCV and DWV-B), queens laid eggs that were primed with an increased number of immune effectors relative to control-injected queens. However, eggs collected from queens in field settings showed no protein, lipid, or metabolite composition changes associated with natural variations in maternal ovary size and virus abundance. The composition of eggs collected from the field, though, does appear to be strongly affected by date, which is consistent with previous reports of seasonal shifts in the adult bee^45,46^ and egg proteome^18^, as well as the seasonal variation in the protein and lipid contained in pollen ^48,49,51^. This work underscores that maternal viral infections in honey bees can indeed trigger a transgenerational effect on the egg proteome, but these effects are likely dependent on the context of the infection, potentially including the route of exposure, timing of exposure, degree of infection, type of virus, the season, nutritional availability, and genetics.

## Methods

### Immune challenge cage experiment

As previously published (see Chapman et al., 2024 ^15^ for full details), young, naturally mated queens were kept in an incubator with approximately 100 attendants in specialized “queen monitoring cages” ^4^ containing plates that enabled them to lay eggs. Queens were allowed to acclimate to the cages and were confirmed to be consistently laying before experimental injection. Queens were then injected on day 6 after introduction to the cages, and observed for 7 more days before sampling. Queens were randomly assigned to each experimental injection group (n = 9 for each group) with the experimenter (Chapman) conducting the experiment blind to group identifiers. Queens were anesthetized with CO_2_ and then injected with a combination of either: (1) infectious BQCV (∼ 1 × 10^8^ copies) and DWV-B (∼ 2 × 10^6^ copies) (live virus group), or (2) the same combination and quantity of virus which had been UV-inactivated (UV-inactivated group), or (3) the same volume of saline (control group). Eggs were collected on day 7 after injection and were no more than 48 hours old, although they were likely mostly younger than this based on anecdotal observations that the queens tended to take 12-24 hours to lay eggs in newly provided, clean egg-laying plates. Blinding was maintained until all data had been collected. Ovaries were dissected immediately upon queen sacrifice and all samples were stored at -70 °C until further processing.

### Queen and egg collection from field sites

Honey bee queens and their eggs were sampled from colonies in the field in 2021. Samples were collected twice from the two British Columbia beekeepers (in Grand Forks and Vernon), first on May 14 & 15 and then again on June 15 & 16, and collected from a Lethbridge, AB beekeeper on June 27 & 28. At each time point, 10 eggs still standing on end (not fallen onto their side, which is indicative of older eggs close to hatching) were collected using a Chinese grafting tool and transferred to Eppendorf tubes. All samples were collected directly onto dry ice and subsequently stored at -70 °C until processing.

### Statistical analysis

The relationship between ovary mass and virus RNA copies in the field queens was analyzed in R (v 4.3.0) using a simple linear model (lm) that was assessed for goodness-of-fit using tools within the DHARMa package ^53^, which utilizes simulation-based residuals to check for typical model misspecification problems such as over-or under-dispersion and residual autocorrelation.

### Proteomic sample preparation and data acquisition

For the ovary samples, one ovary from each queen was homogenized in a 2-mL screw cap tube containing 300 μL of lysis buffer (6 M guanidinium chloride, 100 mM Tris, pH 8.5) and four ceramic beads. For the egg samples, an Eppendorf tube containing 10 eggs was centrifuged for 1 min at 14,000 rcf before 100 μL of lysis buffer was added. A plastic micro pestle was used to grind the eggs until any tissue was no longer visible. The mixture was then transferred to a 2-mL screw cap tube containing 2 ceramic beads along with an additional 50 μL of lysis buffer.

All samples were then homogenized three times for 30 s at 6500 rpm using a Precellys 24 homogenizer (Bertin Instruments) and placed on ice for 5 minutes between each homogenization run. 100 μL of the homogenate was then transferred to a new tube and diluted 1:1 with distilled H_2_O. The protein was precipitated by adding four volumes (800 µL) of ice-cold acetone and incubating overnight at 20 °C. All subsequent sample preparation and mass spectrometry data acquisition was conducted exactly as previously described ^15^. Briefly, the protein was resuspended, quantified using a Bradford assay (PIERCE), and then reduced (dithiothreitol) and alkylated (iodoacetamide). The samples were then digested overnight with trypsin/LysC mixture (Promega) before being acidified with 20% formic acid (to pH < 2.0) and desalted with high-capacity STAGE tips as previously described^54^. A total of 50 ng of digested peptides were injected onto the LC system in randomized order. The digest was separated using NanoElute UHPLC system (Bruker Daltonics) with Aurora Series Gen2 (CSI) analytical column (Ion Opticks, Parkville, Victoria, Australia) coupled to timsTOF Pro (Bruker Daltonics) operated in DIA-PASEF mode for data-independent acquisition scanning.

### Proteomic data processing and analysis

Mass spectrometry data were searched using DIA-NN (v1.8.1) ^55^ as previously described ^15^ using “FASTA digest for library-free search” with “Deep-learning based spectra, RTs and IMs prediction”, allowing for 2 missed cleavages, selecting “Unrelated runs”, setting protein inference to “Protein names from FASTA”, and setting the neural-network classifier to “Double-pass mode”. “N-term M excision” and “C carbamidomethylation” were selected as modifications, and “Use isotopologues”, “MBR”, and “No shared spectra” were all left checked for the algorithm. All other parameters were left set as default. The protein database was the UniProt proteome for *Apis mellifera* (based on the Amel_HAv3.1 genome assembly), plus common honey bee virus and pathogen proteins from UniProt downloaded on February 6, 2023. To this, a comprehensive list of potential protein contaminants was added (Frankenfield et al., 2022).

In R (v 4.3.0), the identified and quantified protein groups from the DIA-NN search were filtered to remove contaminants and proteins identified in fewer than 75% of samples. There were 5,999 protein groups identified before filtering for the eggs from the field survey and 5,574 after. For the eggs from the cage experiment, there were 5,965 protein groups and 5,296 after. For the cage experiment ovaries, there were 5,965 and 5,414 after. The normalized label-free quantification (LFQ) intensity data were then log2 transformed. Differential expression analysis was performed using the limma package in R ^56^. For both the ovaries and the eggs from the cage experiment, group was included as a fixed effect. Contrasts were defined with the control (saline-injected) queens as reference. For assessing the effect of date on the eggs collected from the field survey, the day of collection, the beekeeper, and the mass spectrometry sample injection order were included as fixed effects. The models used to investigate proteins associated with ovary mass or total virus copies (which yielded no significant proteins) included beekeeper and injection order as fixed effects. Empirical Bayes moderation of the standard errors was performed on the resulting outputs, and the false discovery rate was controlled to 5% using the Benjamini–Hochberg correction.

GO term enrichment analysis was performed in ErmineJ ^57^ using the gene score resampling (GSR) method, which tests for enrichment along the continuum of raw p-values as scores rather than comparing enrichments in an arbitrary group of “significant” proteins against all “background” proteins. The false discovery rate for enrichment analysis was controlled to 5% using the Benjamini-Hochberg method.

### Lipidomic and metabolomic sample preparation

Eppendorf tubes containing 10 eggs each were centrifuged for 1 min at 14,000 rcf immediately upon removal from the freezer freezer followed by a two-phase extraction protocol adapted from Matyash et al. (2008)^58^ with some modifications. Briefly, 100 μL of methanol extraction solvent (75% methanol:water, made with 0.01% BHT-methanol and containing standards of methionine-d3, caffeine-^13^C_3_, and ferulic acid-d3 each at 1 ppm and SPLASH LIPIDOMIX (Avanti Polar Lipids, Alabama, USA) at 5 ppm) was added. The eggs were then ground with a mini-pestle to lyse the eggs. The tubes were thoroughly vortexed, and samples were transferred to a 2 mL screw cap tube homogenizer tube. Another 100 μL of extraction solvent was used to rinse the original tube and was also transferred to the homogenization tube to increase yields. 2 ceramic beads were then added and samples were homogenized three times for 30 s at 6500 rpm as described for proteomics sample processing. 500 μL of methyl tert-butyl ether was then added and samples were incubated for 1 hour at room temperature while shaking at 1000 rpm. The mixture was transferred to a new tube and centrifuged at 14,000 rcf for 10 min. 600 μL was transferred to a new tube to which 107 μL of water was added (to a final additional concentration of 15%) before incubating again at room temperature for 10 minutes while shaking at 700 rpm. The samples were then centrifuged for 15 min at 14000 rcf at 4 °C. 300 μL of the upper fraction containing the non-polar (lipid) compounds were transferred to a new tube, while 400 μL of the lower fraction containing the polar (metabolite) compounds were transferred to a different tube. The polar fraction was dried (SpeedVac, Eppendorf) immediately and stored at -70 °C until mass spectrometry analysis. The non-polar fraction was stored in solution at -70 °C and dried immediately before analysis to prevent oxidation.

### Metabolomics LC-MS/MS analysis

Samples were analyzed following previously established protocols in our group^59,60^, with some modifications. The polar fraction was resuspended with 250 µL of 50% aqueous methanol (v/v) and then spun at 16,000 rcf for 10 minutes. After that, 200 µL were transferred into LC vials for LC-MS/MS analysis. A quality control (QC) sample was made by mixing 20 µL from each sample. QCs and blanks were injected approximately every 12 samples. Samples were randomized before injecting 3 µL. The metabolites were separated using an Inertsil Ph-3 UHPLC column (2 µm, 150 × 2.1 mm, GL Sciences) on a Vanquish Horizon UHPLC system (Thermo) connected to an Impact™ II high-resolution mass spectrometer (Bruker Daltonics). The mobile phase consisted of water (A) with 0.1% (v/v) formic acid, and methanol (B) with 0.1% (v/v) formic acid. A multi-step gradient from 5% to 99% of mobile phase B over 18 minutes was used. The column temperature was set at 55°C, the autosampler at 4°C, and the flow rate at 0.3 mL/min. Data-dependent acquisitions were conducted in positive (ESI+) and negative (ESI-) ionization modes. For ESI+, the settings were as follows: a capillary voltage of 4,500 V, nebulizer pressure of 2.0 bar, dry gas flow of 9 L/min, dry gas temperature of 220°C, and a mass scan range of 60– 1,300 m/z with a 0.6-second cycle time. Collision energy was 20 V ramped from 100 to 250% in MS/MS scans. ESI-used a capillary voltage of -3,500 V. Internal calibration was performed using sodium formate (10 µL of 10 mM injected at 0–0.15 min) to ensure high mass accuracy.

### Lipidomics LC-MS/MS analysis

Untargeted lipidomics analysis was conducted as previously described^61,62^ with some modifications. Shortly, analysis was conducted using an Impact II™ high-resolution mass spectrometer (Bruker Daltonics, Bremen, Germany) coupled with an Elute UHPLC system (Bruker Daltonics). Compounds were separated using a multigradient method on an Acquity CSH column (130Å, 1.7 µm, 100 × 2.1 mm, Waters, Milford, MA) equipped with a CSH C18 VanGuard FIT Cartridge (1.7 µm, 2.1 × 5 mm). Mobile phase A consisted of acetonitrile: water (60:40, v/v), and mobile phase B was isopropanol: acetonitrile (90:10, v/v), both containing 0.1% formic acid (v/v) and 10 mM ammonium formate. Separation was achieved using a gradient ranging from 15% to 99% mobile phase B over 17 minutes as follows: 0 min (15% B), 0–2 min (30% B), 2–2.5 min (50% B), 2.5–12 min (80% B), 12–12.5 min (99% B), 12.5–13.5 min (99% B), 13.5–13.7 min (15% B), and 13.7–17 min (15% B). The column was maintained at 65°C with a flow rate of 0.5 mL/min. The injection volume was 2 µL, and the autosampler was kept at 4°C.

A quality control sample was prepared to track instrument performance by pooling 20 µL aliquots from each sample. QC sample was injected every twelve analytical runs. Data-dependent acquisitions were performed in positive ionization mode (ESI+) to obtain precursor and fragment ion information for compound annotation. The mass spectrometer settings were as follows: capillary voltage of 4,500 V, nebulizer gas pressure of 2.0 bar, dry gas flow rate of 9 L/min, dry gas temperature of 220°C, mass scan range of 65–1,700 *m/z*, spectral acquisition rate of 3 Hz, and cycle time of 0.7 s. Collision energy of 20 V was ramped through each MS/MS scan from 100% to 250%. Calibration was carried out by injecting 10 µL of 10 mM sodium formate at the beginning of each run via the 6-port diverter valve.

### Lipidomics and metabolomics data processing

The raw data was processed using Progenesis QI™ software (version V3.0.7600.27622) with METLIN™ plugin (version V1.0.7642.33805, NonLinear Dynamics). This involved peak picking, alignment, deconvolution, normalization, and database searching. Annotations were carried out following established protocols^59,63^ with additional stringency applied. Briefly, only molecular features containing MS/MS data were used to query and match against libraries. Ions detected in quality control (QC) samples were retained if their coefficient of variation (CV) was below 25%. In cases where compounds were detected in both positive and negative ionization modes, the one with the lower CV in QC samples was retained. For level one (L1) annotations, features were matched against the UBC Life Sciences Institute’s in-house spectral library^59^ (Mass Spectrometry Metabolite Library of Standards, MSMLS, IROA Technologies), containing over 500 standards representative of primary metabolism and injected under identical analytical conditions. Level two (L2) annotations were based on screening features against METLIN™^64^, GNPS^65^, HMDB^66^, and LipidBlast^67^. Relative metabolite quantities were calculated by measuring the corresponding peak areas, and normalization was performed using the “all compounds normalization” method, a robust built-in feature of Progenesis QI™ designed for untargeted metabolomics studies.

### Lipidomic and metabolomic data analysis

Differential expression analysis was performed using the limma package in R ^56^ including date of collection, beekeeper and injection order as fixed effects. Metabolite data collected using positive and negative modes was analysed separately to allow for controlling for injection order. The models used to investigate compounds associated with ovary mass or total virus copies (which yielded no significant proteins) included beekeeper and injection order as fixed effects.

### Viral quantification on queens collected from the field

One ovary from each queen was placed in a separate 2 mL screw-cap homogenizer tube containing 500 μL of Trizol reagent and three ceramic beads. The samples were homogenized (Precellys 24 homogenizer, Bertin Instruments) 2 times for 30 s at 6500 rpm and placed on ice for 5 min between runs. The homogenate was then centrifuged for 30 s at 14,000 rcf and all (500 μL) supernatant was transferred to a new tube containing 500 μL of 100% ethanol for nucleic acid precipitation. The rest of the RNA extraction was performed using the Direct-zol RNA Miniprep spin-column kit (Zymo Research), including the DNase I treatment, according to manufacturer instructions. The RNA was eluted in 50 μL of RNase-free water and quantified using a Qubit Fluorometer with the Qubit RNA High Sensitivity assay kit (see Supplementary Data 1 for total RNA extracted from each ovary). A subset of samples were assessed for contaminants using a NanoDrop One spectrophotometer and assessed for RNA integrity using a formaldehyde gel (prepared according to Mansour & Pestov ^68^). The extracted RNA was stored at -70 °C. RNA was diluted further in RNase-free water and 200 ng was used to produce cDNA. Reverse transcription was carried out using the High-Capacity cDNA Reverse Transcription Kit (Applied Biosystems), which utilizes random primers, according to the manufacturer’s instructions. cDNA was kept after synthesis at 4 °C for a maximum of 4 weeks. The RT-qPCR was performed exactly as previously described in Chapman et al. ^15^. Sacbrood virus was also included in the analysis, but was not above the limit for quantification in any of the samples, and was therefore not included.

## Data availability

All proteomics raw data, search results, and search parameters are available on MassIVE (https://massive.ucsd.edu/ProteoSAFe/static/massive.jsp, accession MSV000094114 for the cage experiment which is associated with Figure 1, and accession MSV000095627 for the field data, which is associated with Figures 2 & 3). Sample metadata, including viral abundances, are available in Supplementary Data 1. Processed metabolomics and lipidomics data (along with statistical results and annotation confidence) are provided as Supplementary Data 2 and raw mass spectrometry data are available at Metabolomics Workbench (www.metabolomicsworkbench.org). Source code underlying figures and data analysis are available freely upon request.

## Acknowledgements

Honey bee research in LJF’s group is supported by an NSERC Discovery Grant. Mass spectrometry infrastructure is supported by the Canada Foundation for Innovation, Genome Canada and Genome BC (264PRO, 374PRO) and computational infrastructure is supported by a Digital Research Alliance of Canada Resource Allocation to LJF. A Project *Apis* m. grant to AM, AC, LJF, and DRT funded this research. We want to acknowledge Emily Huxter (Wild Antho) and Liz Huxter (Kettle Valley Queens) for providing the queens from British Columbia; Scott Covey for help developing the RT-qPCR methods; and the UBC proteomics core facility team—Jason Rogalski, Renata Moravcova, and Jeanne Yuan—for running the mass spectrometry samples, instrument maintenance, and technical expertise. Mention of trade names or commercial products in this publication is solely for the purpose of providing specific information and does not imply recommendation or endorsement by the U.S. Department of Agriculture. USDA is an equal opportunity provider and employer.

## Author contributions

AM and AC conceptualized the experiments and analyses with input from LFJ and DRT. AC wrote the first draft of the manuscript, made the figures, and interpreted the data with assistance from AM. LFJ, DRT, AM, RWC, JDF, AM, SERH, KLCW, ZR and KP provided editing assistance. AC conducted the cage experiment. AC and AM collected the queens and eggs from the field, with help from SERH. JF provided the queen monitoring cages and provided guidance on their use. RWC, KP, and ZR produced the purified virus inoculum for use in the immune challenge injections. AC conducted RT-qPCR virus analysis and prepared the ovary samples for proteomic analysis. KW prepared the egg samples for proteomic analysis and assisted AC in preparing the egg samples for lipidomic and metabolomic analysis. AAM conducted the metabolomic and lipidomic analysis. Grants supplied to AC, AM, LJF, and DRT funded the work.

